# Perinatal liver inflammation is associated with persistent elevation of CXCL10 and its canonical receptor CXCR3 on common myeloid progenitors

**DOI:** 10.1101/2024.08.15.607661

**Authors:** Anas Alkhani, Suruthi Baskaran, Abhishek Murti, Blaine Rapp, Claire S Levy, Bruce Wang, Amar Nijagal

## Abstract

Biliary atresia (BA) is a leading cause of liver failure in infants. Despite effective surgical drainage, patients with BA exhibit attenuated immune responses to childhood vaccines, suggesting there are long-lasting alterations to immune function. The perinatal liver is home to hematopoietic stem and progenitor cells (HSPCs) and serves as the epicenter for rapidly progressive and significantly morbid inflammatory diseases like BA. We have previously established the role of neonatal myeloid progenitors in the pathogenesis of perinatal liver inflammation (PLI) and hypothesize that PLI leads to long-term changes to HSPCs in mice that recovered from PLI. To test this hypothesis, we compared the changes that occur to HSPCs and mature myeloid populations in the bone marrow of adult mice during homeostasis and during PLI. Our results demonstrate that HSPCs from animals that recover from PLI (“PLI-recovered”) undergo long-term expansion with a reduced proliferative capacity. Notably, PLI leads to persistent activation of common myeloid progenitors through the involvement of CXCL10 and its canonical receptor, CXCR3. Our data suggests that the CXCR3-CXCL10 axis may mediate the changes in HSPCs that lead to altered immune function observed in BA, providing support for a targetable pathway to mitigate the detrimental long-term immune effects observed in patients with BA.

## Introduction

Biliary atresia (BA) is a rapidly progressive perinatal inflammatory disease of the liver and is the leading indication for liver transplantation in infants and children [1]. BA causes defective biliary drainage leading to liver failure, as well as long-term attenuated immune responses to childhood vaccinations [2–4]. Despite surgery to restore biliary drainage in infancy, the ongoing immune dysfunction in BA leads to inflammation-induced changes to the immune system that persist through childhood. Prevention and treatment of the prolonged attenuated immune response in diseases like BA, requires an understanding of whether perinatal liver inflammation (PLI) in early life leads to long-term changes to immune populations and their progenitors.

The perinatal liver is a particularly distinct environment in being the primary home for hematopoietic stem and progenitor cells (HSPCs) as well as the epicenter for rapidly progressive and significantly morbid inflammatory diseases like BA. HSPCs are central hubs of inflammation [1, 2], and their exposure to inflammatory signals leads to adaptive changes that can alter long-term immunity [3]. Chemokine signaling networks are essential for HSPC retention, recruitment, and differentiation [4, 5]. Tonic inflammatory signals like type I interferons, interleukin 1, and G-CSF, and chemokines like CXCL10, and CXCL12 facilitate migration from the liver to the bone marrow during early life [6, 7]. Inflammation during this transition period perturbs tonic signals, leading to changes to emigrating HSPCs. For example, in utero inflammation activates lymphoid-biased fetal HSPCs, which leads to persistent expansion and hyperactivation of these same fetal-derived lymphoid-biased HSPCs and multipotent progenitors (MPPs) in the postnatal period [8]. Furthermore, inflammation modifies HSPC function by impairing cell renewal and accelerating aging [9]. Notably, elevated levels of CXCL10 in the livers of humans with BA [6] and detected after cholangiocyte injury in congenital hepatic fibrosis [7], suggest a role for chemokine signaling in the pathogenesis of perinatal liver disease.

We recently established a critical role for neonatal liver HSPCs in propagating PLI in neonatal mice [10]. In the current study, we hypothesized that PLI leads to long-term changes to HSPCs in mice that recovered from PLI. To test this hypothesis, we compared the changes that occur to HSPCs and mature myeloid populations in the bone marrow of adult mice during homeostasis and during PLI induced by an infectious model. Our results demonstrate that HSPCs from animals that recover from PLI (“PLI-recovered”) undergo long-term expansion with a reduced proliferative capacity. PLI-recovered HSPCs have increased expression of the proinflammatory cytokine CXCL10 and its canonical receptor, CXCR3. These findings support our hypothesis that PLI leads to long-term changes in the adult bone marrow and suggest the CXCL10-CXCR3 axis is an important one that propagates the perturbations to lifelong immunity observed in patients with perinatal hepatic inflammatory disease.

## Materials and methods

### Mice

8-12-week-old BALB/c mice were obtained from the National Cancer Institute (Wilmington, MA, USA), and received humane care according to the Guide for Care and Use of Laboratory Animals. Mouse experiments were approved by the University of California, San Francisco Institutional Animal Care and Use Committee, and all mice were euthanized according to humane endpoints.

### Postnatal Model of Perinatal Liver Injury

Rhesus Rotavirus (RRV) was grown and titered in Cercopithecus aethiops kidney epithelial (MA104) cells. PLI was induced by intraperitoneal injections (i.p.) of 1.5 x 10^6^ focus forming units (ffu) of RRV within 24 hours of birth. Mice that never showed signs of disease (weight loss, ruffled fur, jaundice) were excluded from the analysis. Controls were injected i.p. with PBS. The liver and bone marrow of RRV– and PBS-injected mice were harvested on day 3 (P3) for flow cytometric experiments. The remaining RRV– and PBS-injected mice were maintained for 8-12 weeks in a BSLII facility until they recovered from the initial infection (PLI-recovered mice and PBS-injected controls). The liver, spleen, bone marrow, and serum were harvested from these mice for flow-cytometric analysis, bulk RNA sequencing, and cytokine analysis.

### Creation of Single Cell Suspension

Bone marrow: Femurs and tibias of adult mice were harvested, cleaned, and flushed with 1ml of sterile cold PBS, and underwent mechanical dissociation and filtration through a 100µm strainer for further processing. For P3 mice, the femur, tibia, pelvis, and ischial spines were harvested and crushed using a mortar and pestle. The tissue was then washed with cold saline and filtered through a 100µm strainer.

Liver: Liver was harvested and washed three times with cold sterile PBS, minced, and then incubated in prewarmed 0.2 mg/ml C. histolyticum collagenase (Sigma Aldrich, St. Louis, MO, USA C5138) RPMI solution for 30 minutes rotating at room temperature (collagenase was only used for adult livers). The solution was then mechanically dissociated to ensure complete digestion of tissue and filtered through a 100µm strainer for further processing. For P3 livers, mechanical dissociation alone was used to create a single cell suspension.

Spleen: Spleen was harvested and washed with cold sterile PBS followed by mechanical dissociation through 100µm strainer for further processing.

Serum: Serum was collected under sterile conditions and isoflurane anesthesia using cardiac puncture according to IACUC approved protocols.

### Flow Cytometry and Sorting

Single-cell suspensions from the bone marrow were divided into two fractions. One fraction along with single-cell suspensions from the liver and spleen were stained for surface markers to identify mature myeloid cells (**Supplementary Table S1**). To isolate HSPCs, the other fraction was depleted of lineage-positive (Lin+) cells using a Direct Lineage Cell Depletion Kit (Miltenyi Biotec, Cambridge, MA, USA, 130-110-470) containing antibodies to CD5, CD45R, CD11b, Gr-1, 7-4, Ter-119 and stained for the following HSPCs: long-term hematopoietic stem cells (HSC^LT^), common myeloid progenitors (CMP^+^ and CMP^−^), terminal myeloid progenitors (TMPs; megakaryocytic erythroid progenitors, MEP; monocytic-dendritic progenitors, MDP; granulocytic-monocytic progenitors, GMP; granulocytic progenitors, GP; monocytic and common monocytic progenitors, MP) using cell surface markers (**Supplementary Table S2**). Flow cytometry was performed on an LSR Fortessa X20 (BD Biosciences, San Jose, CA, USA). Flow sorting for CMPs (both CMP^+^ and CMP^−^) was performed using a BD FACSAria2 SORP (BD Biosciences, San Jose, CA, USA) and data were analyzed using FlowJo 10.10.0 (Ashland, OR, USA).

### Luminex ProcartaPlex Analysis

Luminex ProcartaPlex assay was performed using ProcartaPlex™ Mouse Cytokine & Chemokine Convenience Panel 1A 36-Plex (ThermoFisher Scientific, EPXR360-26092-901). The serum from PLI-recovered mice and PBS-injected controls were centrifuged at 10,000g. 4-fold serial dilutions of the standard mix were also prepared. The capture beads were added to a 96-well plate followed by 25µL of the standard or serum samples in duplicates along with 25µL of universal assay buffer. This was incubated for 2 hours and then washed. The plate was then incubated with 25µL of detection antibody mix, washed and incubated again with 50µL Streptavidin-PE. The beads were then suspended in 120µL of reading buffer. The plate was read using Bio-Rad Bio-Plex 200 self-use analyzer and levels of 36 cytokines were recorded. The final concentrations of each cytokine were determined based on the standard curves using the Invitrogen™ ProcartaPlex™ Analysis App.

### Bulk-RNA Sequencing Analysis

The HSPCs from the bone marrow of adult PLI-recovered and PBS-injected mice were stained with surface antibodies and flow-sorted to collect CMP^+^ and CMP^−^ fractions in RNAlater solution (Sigma-Aldrich, St. Louis, MO, USA R0901). They were then analyzed for bulk RNA sequencing. MedGenome, Inc. (Foster City, CA, USA) performed quality control analysis, RNA extraction, and raw data generation. Using an unbiased approach, the raw data was analyzed for differential gene expression using the DESeq2 package. The raw fastq files were aligned to the GRCm39 mouse genome using the STAR aligner. The count matrices were generated using featureCounts. The final gene list was compiled using a cut-off for log_2_ fold change of <-0.5 and >0.5. Differentially expressed genes were categorized into functionally-related groups using the Database for Annotation, Visualization and Integrated Discovery (DAVID).

### Data Analysis and Statistics

All graphs and statistics were generated using GraphPad Prism 9.3.1 (San Diego, CA, USA). Individual proportions of HSPCs were calculated based on absolute cell counts (ACC) as either a percentage (%) of the total lineage-negative progenitor compartment (Lin^−ve^ cells) or as a fraction of total HSC^LT^ and downstream myeloid progenitors (CMPs and TMPs). Mature myeloid cell proportions were calculated as a percentage of CD45^+^ leukocytes based on ACC. The Mann– Whitney test was used to compare the proportion and ACC of HSPCs, mature myeloid cells, and cytokine levels. Unpaired t-tests were used to compare CXCR3 expression of neonatal HSPCs after PBS and RRV infection. A p-value of <0.05 was considered significant. Error bars represent mean ± standard error mean (SEM). All authors had access to the study data and reviewed and approved the final manuscript.

## Results

### PLI increases the number of myeloid progenitors in the bone marrow of PLI-recovered adult mice

To test whether PLI results in long-term changes to HSPCs, we injected neonatal mice with 1.5×10^6^ffu of RRV within 24 hours of birth. As expected, RRV infection led to bile duct injury and periportal liver inflammation, an injury pattern that resembles human BA [11]. RRV-induced PLI progressed between days 3-14. Of the 45 Balb/c mice that were injected, 64% died (n=16) and 36% recovered from PLI (“PLI-recovered”) (**Fig. 1a**), findings comparable to our published results describing the mortality associated with RRV-mediated PLI (5).

**Figure 1.**
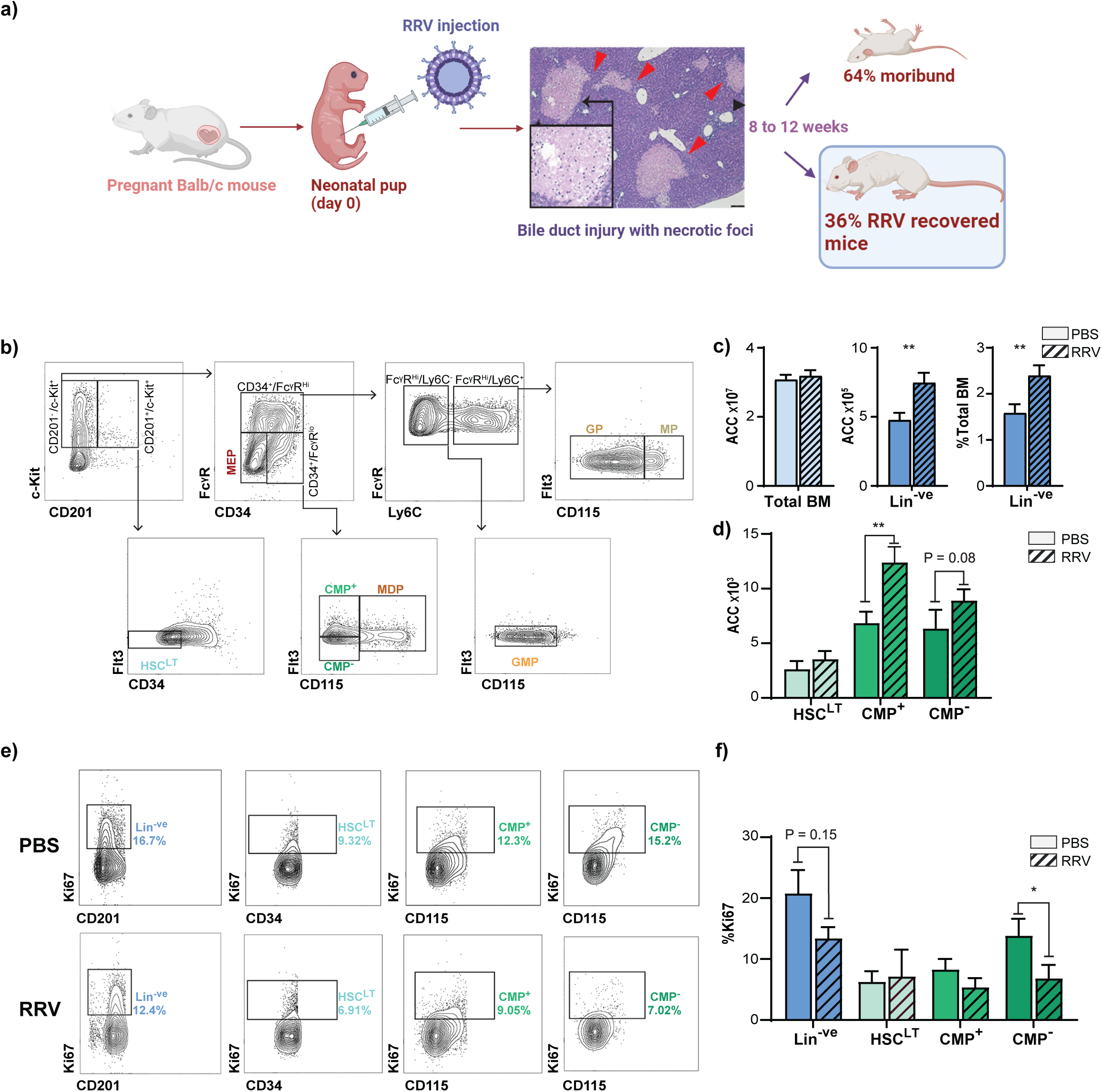
PLI results in expansion and reduced proliferation of lineage negative cells (Lin-ve) in the bone marrow of PLI-recovered adult animals. a) Schematic showing generation of PLI-recovered adult mice. b) Plots demonstrating flow-cytometric gating strategy of hematopoietic stem and progenitor cells (HSPCs) in adult mice bone marrow (BM) including long-term hematopoietic stem cells (HSC^LT^), common myeloid progenitors (CMP^+^ and CMP^−^), megakaryocytic-erythrocytic progenitors (MEP), monocytic-dendritic progenitors (MDP), granulocytic-monocytic progenitors (GMP), monocytic progenitors (MP), and granulocytic progenitors (GP). c) Bar graphs (left-right) showing no change in total bone marrow absolute cell count (ACC), significant increase in Lin^−ve^ ACC (p-value = 0.004+/− SEM) and percentage (p-value = 0.002+/− SEM) in PLI-recovered animals compared to (c.t) PBS-injected controls. d) Bar graph showing quantification of ACC in HSC^LT^, CMP^+^ and CMP^−^ with significant increase in CMP^+^ in PLI-recovered animals (p-value = 0.005+/− SEM) c.t. PBS-injected controls. Flow cytometric plots (e) and bar-graph (f) showing Ki67% expression in Lin^−ve^, HSC^LT^, CMP^+^ and CMP^−^ populations with significantly lower Ki67% expression reflected in CMP^−^ (p-value= 0.04+/− SEM) in PLI-recovered animals (n=22) c.t. PBS-injected controls (n=11). p-value * < 0.05; ** < 0.01. Error bars represent mean +/− SEM.

Since our previous studies implicated liver HSPCs in the pathogenesis of PLI in neonatal mice, we were interested in whether PLI also exerted long-term effects on HSPCs in PLI-recovered mice. We used flow cyometry to quantify hematopoietic stem and progenitor cells (HSPCs – includes HSC^LT^, CMP^+^ (Flt3+ common myeloid progenitors), CMP^−^ (Flt3-common myeloid progenitors), GMP (granulocytic-monocytic progenitors), MDP (monocytic-dendritic progenitors), MEP (megakaryocytic-erythroid progenitors), GP (granulocytic progenitors), MP (monocytic and common monocytic progenitors)) using flow cytometry in the bone marrow of PLI-recovered adult mice (**Fig. 1b**) and compared the absolute cell number and proportion of HSC^LT^ (Sca1^+^c-Kit^+^CD34^−^Flt3^−^), CMP^+^ (Sca1^−^c-Kit^+^CD34^+^FcᵞR^lo^Flt3^+^CD115^lo^), and CMP^−^ (Sca1^−^c-Kit^+^CD34^+^ FcᵞR^lo^Flt3^−^CD115^lo^) to that of PBS-injected controls. The livers of adult PLI-recovered mice had a limited number of HSPCs (*data not shown*), leading us to focus our analysis on HSPCs in the bone marrow of PLI-recovered mice. The total number of cells in the bone marrow was unchanged in PLI-recovered mice compared to controls (**Fig. 1c**). However, a significantly higher number and proportion of lineage-negative (Lin^−ve^) cells (bone marrow progenitors after depletion using antibodies to CD5, CD45R, CD11b, Gr-1, 7-4, Ter-119) were found in PLI-recovered mice than in controls (**Fig. 1c**). There was no difference in the absolute cell counts of HSC^LT^ when comparing PLI-recovered mice and controls. We did, however, observe significantly higher counts of CMP^+^, and higher (but not significant) counts of CMP^−^ (**Fig 1d**). The two groups had similar proportions of HSC^LT^, CMP^+^, CMP^−,^ and terminal myeloid progenitors such as GMP (Sca1^−^c-Kit^+^CD34^+^ FcᵞR^Hi^Flt3^−^ CD115^lo^), MEP (Sca1^−^c-Kit^+^CD34^−^FcᵞR^lo^), MDP (Sca1^−^c-Kit^+^CD34^−^FcᵞR^lo^Flt3^+^CD115^Hi^), GP (Sca1^−^c-Kit^+^CD34^+^FcᵞR^Hi^Flt3^−^Ly6C^+^CD115^lo^), and MP (Sca1^−^c-Kit^+^CD34^+^ FcᵞR^Hi^ Flt3^−^ Ly6C^+^ CD115^Hi^) (**Supp.** Fig. 1a). The two groups also had similar absolute cell counts of terminal myeloid progenitors including GMP, MEP, MDP, GP but PLI-recovered mice had a significantly higher MP count in the bone marrow (**Supp.** Fig. 1b). These findings indicate that PLI leads to a significant expansion of bone marrow Lin^−ve^ cells, CMP^+^ HSPCs and MPs in adult, PLI-recovered mice.

Next, we sought to determine whether expansion of the Lin^−ve^ fraction in the bone marrow of PLI-recovered mice was due to increased proliferation. For this, we measured Ki67 expression in Lin^−ve^, HSC^LT^, CMP^+^, and CMP^−^ by flow cytometry (**Fig. 1e**). Surprisingly, Ki67 expression decreased in the Lin^−ve^ (p=0.15) and CMP^−^ (p=0.04) populations in the bone marrow of PLI-recovered mice, indicating diminished proliferation of these populations. Ki67 expression was similar among HSC^LT^ (Sca-1^+^ c-Kit^+^ CD34^−^ Flt3^−^) and CMP^+^ fractions (**Fig. 1f**), and downstream myeloid progenitors (GMP, MEP, MDP, GP, MP) (**Supp.** Fig. 1c). These findings indicate that the expanded Lin^−ve^ cells and CMPs are quiescent in the bone marrow of PLI-recovered mice.

The increase in proportion of CMPs in PLI-recovered mice led us to ask whether downstream myeloid cells were altered in these mice. To answer this question, we used flow cytometry to evaluate mature myeloid populations including neutrophils, Ly6c^Hi^ and Ly6c^Lo^ monocytes, monocyte-derived macrophages and tissue macrophages in the bone marrow, liver, and spleen of PLI-recovered mice using flow cytometry (**Supp.** Fig. 2a). We found similar percentages of total CD45^+^ hematopoietic cells and mature myeloid populations in the bone marrow (**Supp.** Fig. 2b), liver (**Supp.** Fig. 2d), and spleen (**Supp.** Fig. 2f) of PLI-recovered and PBS-injected mice. In the two groups, the absolute numbers of mature myeloid populations were also similar in the bone marrow (**Supp.** Fig. 2c), liver (**Supp.** Fig. 2e), and spleen (**Supp.** Fig. 2g). Collectively, these results indicate that PLI causes expansion of a quiescent population of HSPC compartment without resulting in quantitative differences in mature myeloid populations in PLI-recovered mice.

### PLI leads to upregulation of pro-inflammatory genes in CMPs

Inflammatory signals alter the transcriptional and epigenetic memory of HSPCs [12]. During perinatal life, maternal inflammatory signals lead to transcriptional changes to fetal liver HSPCs [8], resulting in the persistent expansion of developmentally restricted HSPCs in adult bone marrow [8]. To determine whether these transcriptional changes to bone marrow HSPCs occur after recovery from PLI, we performed bulk RNA sequencing of CMPs (CMP^+^ and CMP^−^) from the bone marrow of PLI-recovered mice and PBS-injected controls (**Fig. 2a**). Compared to CMPs from controls, CMPs from PLI-recovered adult mice had increased expression of genes related to basic cellular pathways such as cell cycle (*Lipg, Il18bp, Cd74, Hba-a1, Nod1*), metabolism (*Dhrs3, Degs2, Lipg, Ldlr, Hba-a1, Hba-a2, Hbb-bs, Syde1, Reck, Nod1*), and membrane function (*Lysmd4, Dhrs3, Disp2, Parm1, Sapcd2*) and had decreased expression of genes related to protein synthesis and stress response (*Gzmb, Hspa8, Hsph1, Thsd4, Pde3b*) (**Fig. 2b-c**). In particular, the CMPs from PLI-recovered mice showed upregulation of genes related to immune function such as macrophage cytokine production (*Cd74, Nod1*), dendritic cell antigen processing (*Cd74, Nod1*), regulation of the pro-inflammatory cytokines IL-6 and IL-8 (*Il18bp, Cd74, Nod1*), and chemokine receptor function (*Cxcr3*) (**Fig. 2b-c**). These findings suggest that pro-inflammatory pathways in CMPs are persistently activated in PLI-recovered mice.

**Figure 2.**
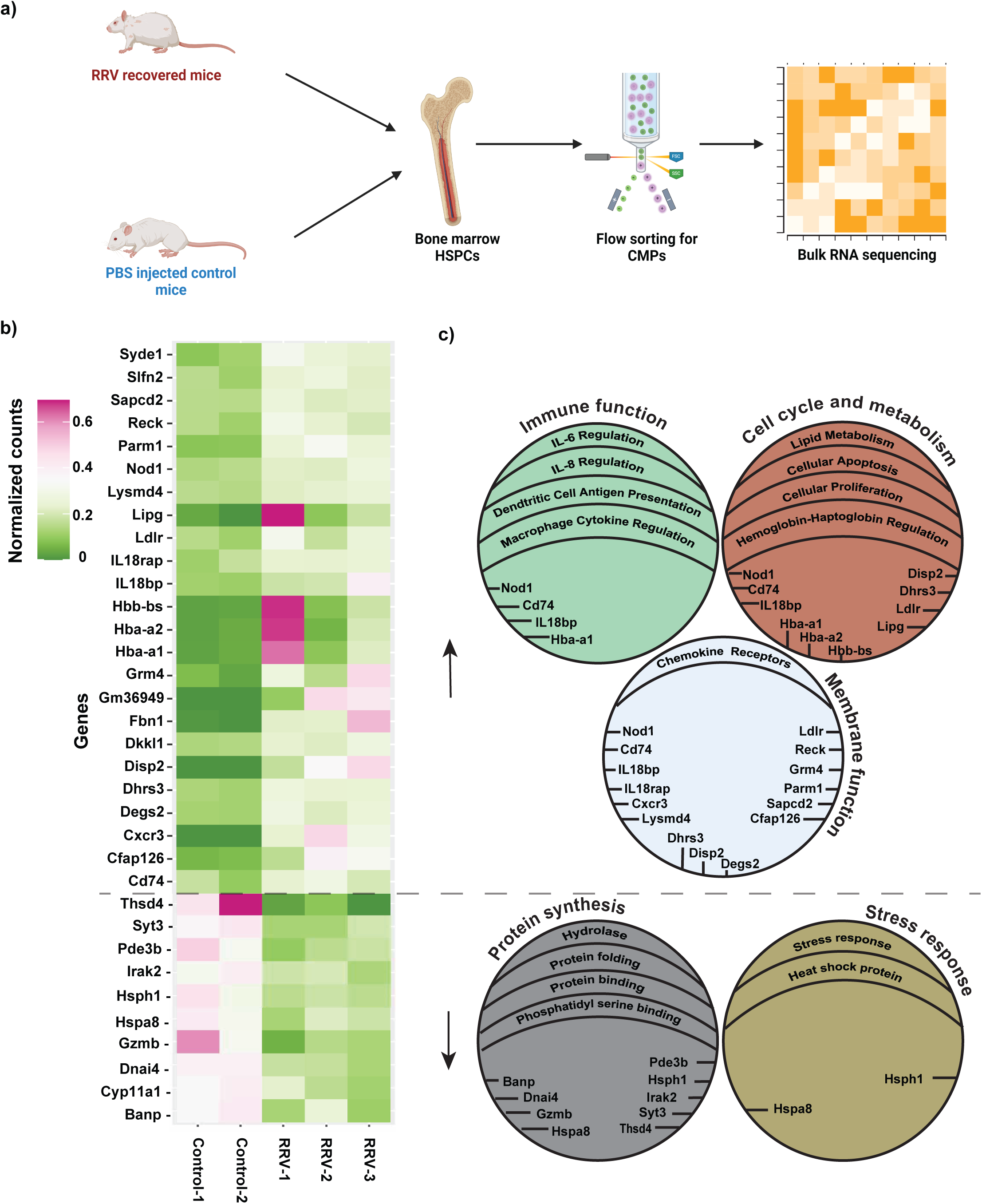
PLI alters gene expression of CMPs in PLI-recovered adult mice. a) Schematic showing the experimental protocol for bulk RNA sequencing of CMPs from PLI-recovered mice. b) Heatmap showing differential gene expression including upregulation of proinflammatory genes such as *IL18rap, IL18bp, Cxcr3, Nod1* in CMPs sorted from PLI-recovered animals bone marrow (n=3) c.t. PBS-injected controls (n=2). c) Venn diagram illustrating functional annotation and clustering of upregulated (above dotted line) and downregulated (below dotted line) genes in CMPs of PLI-recovered mice.

### PLI leads to persistent elevation of CXCL10 and CXCR3

According to our findings, PLI-recovered mice have persistent expansion and transcriptional activation of pro-inflammatory genes of adult bone marrow HSPCs. To ascertain whether the persistent changes in HSPCs could be explained by changes in systemic immune signaling, we measured cytokines and chemokine levels in the serum of PLI-recovered animals and PBS-injected controls. We found that the levels of the pro-inflammatory cytokines CXCL5, CCL11, IL5, IL6, IL27, and CCL7 tended to be higher in the PLI-recovered mice, but were not significantly different from those in controls. PLI-recovered mice also tended to have lower levels of the anti-inflammatory cytokine IL28 in PLI-recovered animals (**Fig. 3a**).

**Figure 3.**
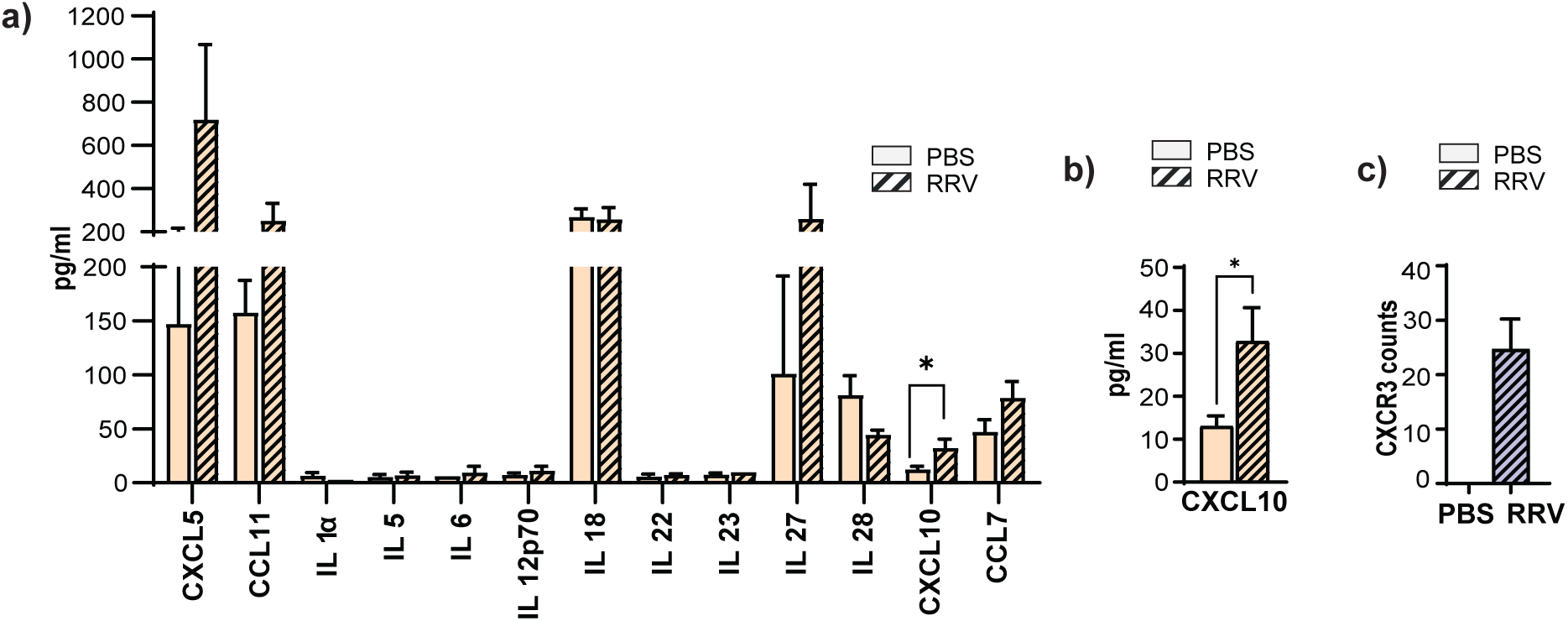
PLI leads to increased activity of the CXCR3-CXCL10 axis in PLI-recovered adult mice. a) Bar graph representing elevated levels of pro-inflammatory cytokines CXCL5, CCL11, IL27, CCL7 and lower levels of the anti-inflammatory cytokine IL28, without reaching significance in PLI-recovered mice (n=4) c.t PBS-injected controls (n=5). b) Bar graph depicting significantly higher levels of the pro-inflammatory cytokine CXCL10 in PLI-recovered animals c) t. controls (p-value = 0.03+/− SEM). c) Bar graph representing increased normalized counts of CXCR3 gene in the CMPs from bone marrow of PLI-recovered mice c.t controls. p-value * < 0.05, ** < 0.01. Error bars represent mean +/− SEM.

PLI did lead to significantly higher levels of CXCL10, which persisted (**Fig. 3b**) even without ongoing liver inflammation (*data not shown*). CXCL10 and its canonical receptor CXCR3 play a role in HSPC migration, retention, and differentiation [7]. To determine whether changes in CXCL10 were also associated with changes to its canonical receptor CXCR3 we examined levels of CXCR3 transcripts. PLI-recovered mice had high levels of these transcripts in CMPs whereas CXCR3 mRNA was not detected in PBS-injected controls (**Fig. 3c**). These findings confirm that PLI is associated with persistently increased serum CXCL10 and CXCR3 expression on adult CMPs implicating this pathway as a potential mechanism by which PLI leads to long-term effects on HSPCs.

### PLI results in increased expression of CXCR3 in hepatic CMPs early in the course of PLI

Our next step was to determine whether CXCR3 expression on HSPCs increases during the early phases of PLI in neonatal mice. For this, we measured CXCR3 expression in bone marrow and liver three days after RRV injection (**Fig. 4a**). We found that in mice with PLI, CXCR3 expression increased significantly in CMP^+^ HSPCs in the bone marrow and liver (**Fig. 4b**) compared to controls. In neonatal mice with PLI, CXCR3 expression on liver CMP^+^ HSPCs tended to be increased compared to that in the bone marrow (**Fig. 4b**). CXCR3 also tended to be increased in bone marrow CMP^−^ HSPCs in mice with PLI compared to controls (**Fig. 4c**), and in liver CMP^−^ of mice with PLI compared to liver of controls (**Fig. 4c**). Finally, in mice with PLI, CXCR3 expression was significantly higher in the CMP^−^ population in the liver than in the bone marrow (**Fig. 4c**). Collectively, our findings indicate that PLI increases CXCR3 expression in hepatic CMPs at the time of disease initiation and this difference is more pronounced in the liver than in the bone marrow.

**Figure 4.**
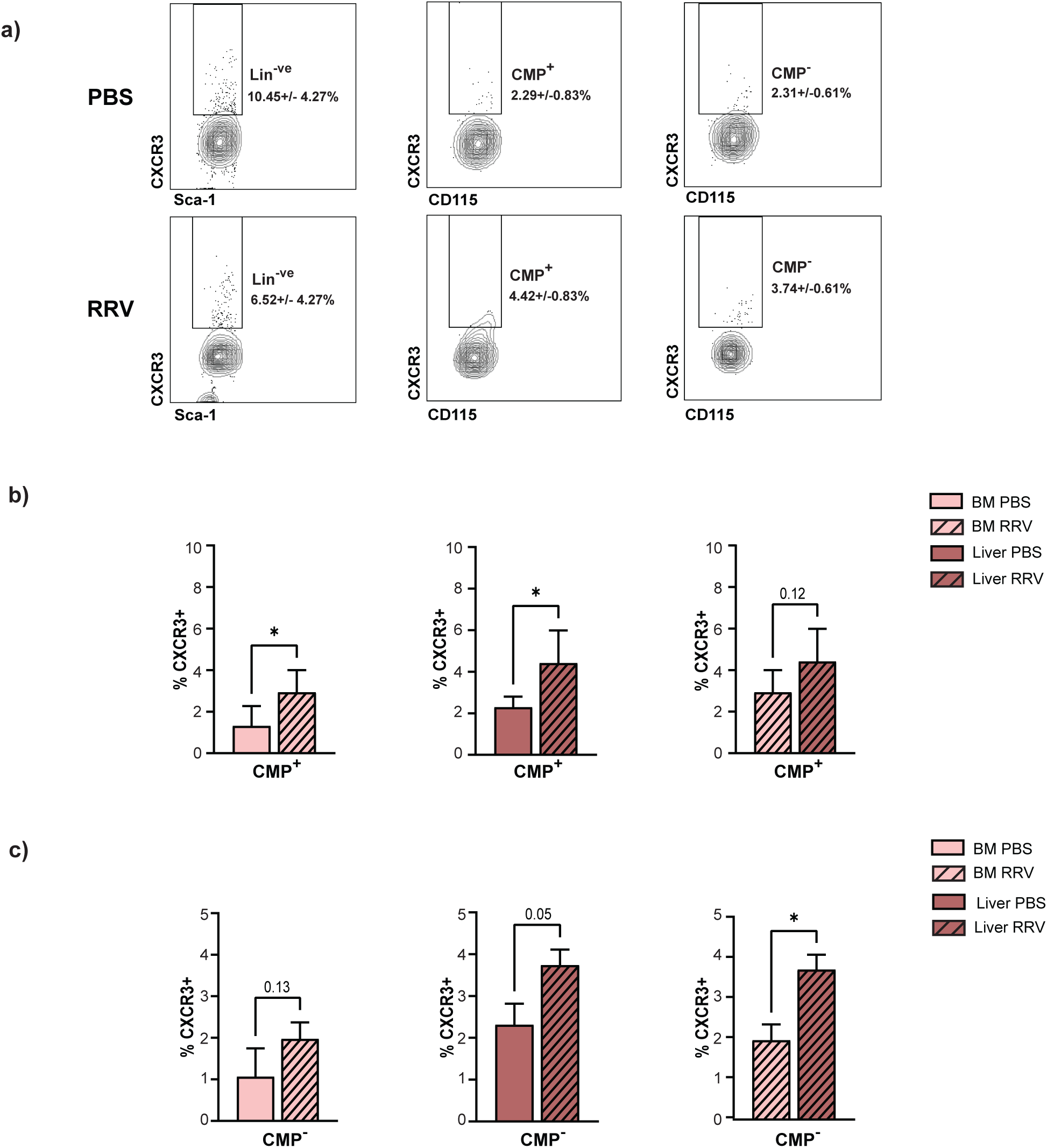
PLI leads to upregulation of CXCR3 in CMPs from neonatal mice with PLI. a) Plots demonstrating flow cytometric gating strategy for CXCR3 expression of HSPCs in the liver of PBS-injected (n=4) and PLI-induced (n=5) neonatal mice. b) Bar graphs depicting CXCR3 expression in the CMP^+^ population of neonatal animals, upregulation of CXCR3 expression in bone marrow of neonatal animals with PLI c.t bone marrow of PBS-injected controls (p-value = 0.049 +/− SEM), and in liver of neonatal animals with PLI c.t liver of PBS-injected controls (p-value = 0.037 +/− SEM). c) Bar graphs depicting CXCR3 expression in the CMP^−^ population of neonatal animals, upregulation of CXCR3 expression in liver c.t bone marrow of neonatal animals with PLI (p-value = 0.012 +/− SEM). p-value * < 0.05; ** < 0.01. Error bars represent mean +/− SEM.

## Discussion

In this study, we hypothesized that PLI leads to long term changes to hematopoietic progenitors in adult mice that have recovered from PLI. Our findings indicate that PLI: (1) results in expansion of a quiescent population of Lin^−ve^ and CMPs cells in the bone marrow of these mice, (2) sustains expression of several pro-inflammatory genes in CMPs in the absence of ongoing liver inflammation, (3) causes increased expression of CXCR3 on CMPs and its ligand, CXCL10, in the serum of PLI-recovered mice, (4) leads to increased CXCR3 expression on CMP^+^ and CMP^−^ populations of the neonatal liver, and (5) may lead to long-term expansion of quiescent CMPs in the adult bone marrow through the persistent activation of the pro-inflammatory CXCR3-CXCL10 axis. These findings suggest that PLI leads to activation of CXCR3 in neonatal liver CMPs (through possible release of CXCL10 following cholangiocyte injury) which could lead to retention and transcriptional changes to CMPs. These maladapted CMPs, upon migration to adult bone marrow, result in long-term expansion of quiescent CMPs that have persistent pro-inflammatory CXCR3-CXCL10 signaling **(Figure 5)**.

**Figure 5.**
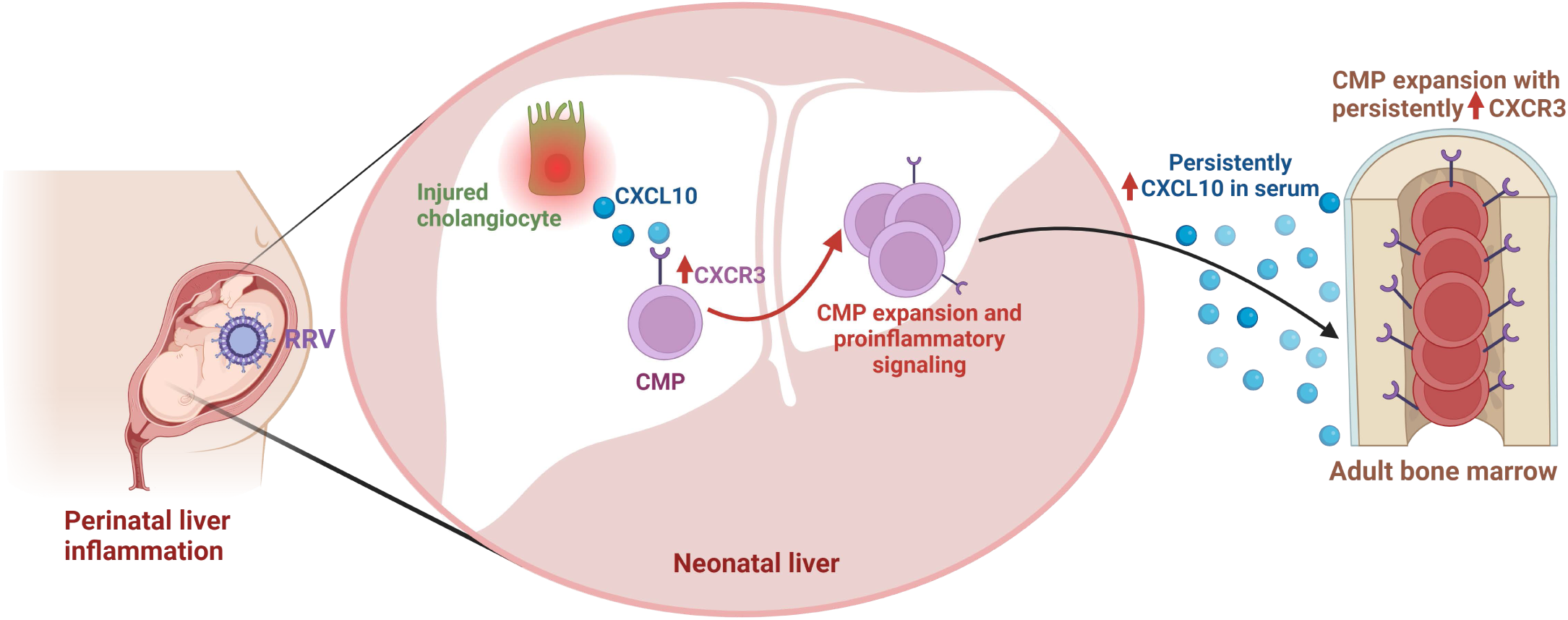
A proposed model showing the role of the CXCR3-CXCL10 axis in propagating long-term effects of PLI. PLI induced by Rhesus Rotavirus (RRV) causes injury to cholangiocytes in the neonatal liver. This could lead to a release of the pro-inflammatory cytokine CXCL10 which signals through its canonical receptor CXCR3 on neonatal liver CMPs. This interaction could in turn lead to the expansion of the pro-inflammatory neonatal liver CMPs which eventually migrate to the bone marrow in adult life. These adult bone marrow CMPs expand and retain the pro-inflammatory changes including increased CXCR3 expression and persistently high CXCL10 levels in the adult serum.

We previously established that myeloid progenitors in the neonatal liver expand and propagate the effects of PLI, and their depletion improves the survival of affected mice [10]. Our current findings demonstrate that the inflammatory insult continues to exert effects on HSPCs by expanding Lin-ve cells and CMPs, indicating that early-life liver inflammation can cause lasting changes to HSPCs that persist in adult animals. These findings corroborate the known link between perinatal inflammatory exposure and long-term modifications to postnatal hematopoietic stem cell activity [8]. Our unexpected Ki67 results showing that the total HSPCs and common myeloid progenitors are quiescent despite an expanded HSPC population. This leads us to speculate whether the adult bone marrow HSPCs were modified by perinatal liver inflammatory signals to evade apoptosis. Such evasion of apoptosis by chronic inflammation-primed HSPCs reportedly contributes to clonal expansion of HSPCs [13]. When populations downstream of HSPCs are studied, the lack of changes in the mature myeloid compartment of PLI-recovered animals indicates the possibility that the changes to HSPCs do not translate to ongoing changes to their differentiation capacity. Despite the lack of change to differentiation capacity, our data supports the conclusion that the transcriptomes of PLI-exposed HSPCs are altered and that these changes may affect HSPC functions other than differentiation. Thus, our results support the idea that perinatal liver inflammatory insult plays a crucial role in the persistence of an expanded, quiescent HSPC subset in the adult bone marrow.

Our findings from bulk RNA sequencing of CMPs and cytokine analysis from PLI-recovered mice further bolster the link between inflammatory signals and long-term changes to HSPC biology [2, 14–16]. These PLI-primed CMPs demonstrate an upregulation of pro-inflammatory genes such as IL-6, IL-8 and the chemokine receptor CXCR3. IL-6 contributes to the proliferation and myeloid differentiation of HSPCs [1], which further correlate with the long-term expansion of HSPCs and common myeloid progenitors observed after recovery from PLI. CXCR3 is commonly known to be located on memory and activated T lymphocytes [17], macrophages [18], and natural killer cells [19]. However, CXCR3 receptors induced by GM-CSF on CD34+ HSPCs were recently found to be crucial for HSPC chemotaxis, differentiation, and maturation [20]. CXCR3 exerts its action by binding the chemokine CXCL10 to affect the migration, differentiation, and activation of hematopoietic and immune cells [21]. Notably, the levels of pro-inflammatory chemokine CXCL10 correlate with the severity of perinatal liver inflammatory diseases like BA [22]. Our findings of CXCR3 upregulation in PLI-primed neonatal liver HSPCs and the activation of the CXCR3-CXCL10 axis in adult mice that recovered from PLI point to the pivotal role of this pro-inflammatory axis in modifying long-term HSPC biology. These changes to HSPCs could potentially contribute to the prolonged immune dysfunction observed in BA and other perinatal liver inflammatory diseases [23–25]. Notably, chronic inflammatory diseases such as atherosclerosis demonstrate similar maladaptive immune responses following primary exposure to an inflammatory stimulus [26].

Our study supports the idea that manipulation of HSPCs could not only treat the immediate effects of PLI but also eliminate the long-term immune dysfunction that these patients with PLI experience. BA, for example, is a perinatal liver inflammatory disease that leads to progressive liver failure despite surgical procedures like Kasai portoenterostomy to re-establish bile flow [27]. A phase I trial of HSPC manipulation using granulocyte colony-stimulating factor in post-Kasai BA patients has been found to improve short-term biliary drainage [28]. The possible role of the CXCR3-CXCL10 axis in causing long-term changes to HSPCs suggests that inhibition of the CXCR3-CXCL10 axis on HSPCs could prevent the immune dysfunction patients experience. Such inhibition of CXCR3-CXCL10 to treat diseases is not uncommon and has been studied extensively in the management of cancers [21, 29, 30], cardiovascular diseases [31] and Listeria monocytogenes-induced fetal wastage [32]. However, essential prerequisites before therapeutic testing include future mechanistic studies to prove the causative role of the CXCR3-CXCL10 axis in causing maladaptive changes to neonatal liver HSPCs and tracing studies to see if maladapted neonatal liver HSPCs eventually migrate to the adult bone marrow.

In conclusion, our study demonstrates that PLI leads to long-term expansion of a quiescent bone marrow HSPC subset through possible activation of the pro-inflammatory CXCR3-CXCL10 axis. Further studies are required to prove the mechanistic role of CXCR3-CXCL10 signaling in modifying HSPCs and the effectiveness of antagonists to this axis in the management of perinatal liver inflammatory diseases. Manipulating HSPCs via the CXCR3-CXCL10 axis may have a role in preventing and treating both acute PLI as well as long-term immune dysfunction in patients with biliary atresia.

## Acknowledgements

The authors would like to thank Dr. Henry Greenberg (Stanford University, CA) for providing MA104 cells and Rhesus rotavirus. The authors would also like to acknowledge Pamela Derish (UCSF Department of Surgery) for critical review of the manuscript. Funding was provided by the American Pediatric Surgical Association Foundation Jay Grosfeld, MD Scholar Award (AN), an American College of Surgeons Faculty Research Fellowship (AN), a UCSF Liver Center Pilot Award (NIH P30 DK026743, AN), the UCSF Society of Hellman Fellows Award (AN), the UCSF Parnassus Flow Cytometry Core (DRC Center Grant NIH P30 DK063720), and core resources of the UCSF Liver Center (P30 DK026743).

