## Supplementary materials for "Perinatal liver inflammation is associated with persistent elevation of CXCL10 and its canonical receptor CXCR3 on common myeloid progenitors"

**Supplementary material:**

**Supplementary Table S1. Staining panel for mature myeloid cells**

| Antibody | Clone | Manufacturer | Catalog number |
|---|---|---|---|
| MHCII | M5/114.15.2 | eBioscience, Waltham, MA | 48-5321-82 |
| Cd11b | M1/70 | eBioscience, Waltham, MA | 47-0112-82 |
| Cd45 | 30-F11 | eBioscience, Waltham, MA | 56-0451-82 |
| Ghost | Not relevant | Tonbo Biosciences, San Diego, CA | 13-0870-T100 |
| Ly6C | AL-21 | BD Biosciences, Franklin Lakes, NJ | 563011 |
| Ly6G | 1A8 | BD Biosciences, Franklin Lakes, NJ | 560601 |
| Fc Block Cd16/Cd32 | 2.4G2 | BD Biosciences, Franklin Lakes, NJ | 553142 |
| Cd11c | N418 | Biolegend, San Diego, CA | 117339 |
| Cd64 | X54-5/7.1 | Biolegend, San Diego, CA | 139323 |

**Supplementary Table S2. Staining panel for hematopoietic stem- and progenitor cells (HSPCs)**

| Antibody | Clone | Manufacturer | Catalog number |
|---|---|---|---|
| c-kit CD117 | 2B8 | Biolegend, San Diego, CA | 105820 |
| Sca-1 Ly6A/E | D7 | Biolegend, San Diego, CA | 108114 |
| FcγR Cd16/Cd32 | 2.4G2 | BD Biosciences, Franklin Lakes, NJ | 553142 |
| LY6C | HK1.4 | Biolegend, San Diego, CA | 128012 |
| CSF-1R CD115 | AFS98 | Biolegend, San Diego, CA | 135506 |
| Flt3 CD135 | A2F10.1 | BD Biosciences, Franklin Lakes, NJ | 560718 |
| CD34 | RAM34 | BD Biosciences, Franklin Lakes, NJ | 553733 |
| CXCR3 | CXCR3-173 | Biolegend, San Diego, CA | 126531 |
| Ki67 | B56 | BD Biosciences, Franklin Lakes, NJ | 561165 |

**Supplementary Figure Legends**

**Supplementary Figure 1. Perinatal liver inflammation leads to significantly increased number of myeloid progenitors with no changes in the percentage and proliferative capacity of downstream progenitor populations in PLI-recovered adult mice.** a) Bar graphs showing no change in the fraction of $HSC^{LT}$, $CMP^{+}$, $CMP^{-}$, GMP, MEP, GP and MP in the $Lin^{-ve}$ compartment in the bone marrow of PLI-recovered animals (n=22) c.t. PBS-injected controls (n=11). b) Bar graph showing quantification of ACC in GMP, MEP, GP, and MP with significant increase in MP in PLI-recovered animals (n=22) (p-value = 0.009+/- SEM) c.t. PBS-injected controls (n=11). c) Bar graph showing no change in Ki67% expression in GMP, MEP, GP and MP populations in PLI-recovered animals (n=22) c.t. PBS-injected controls (n=11). Error bars represent mean +/- SEM.

**Supplementary Figure 2. Perinatal liver inflammation does not lead to any quantitative changes in the mature myeloid compartment of PLI-recovered adult mice.** a) Plots demonstrating flow cytometric gating strategy of mature myeloid populations in adult mice bone marrow. Bar graphs (left-right) showing quantification of percentage CD45+ cells, percentage and ACC of mature myeloid populations in bone marrow (b and c), liver (d and e), and spleen (f and g) of PLI-recovered animals (BM: n=13, Liver: n=5, Spleen: n=14) c.t. PBS-injected controls (BM: n=12, Liver: n=6, Spleen: n=12). Error bars represent mean +/- SEM.

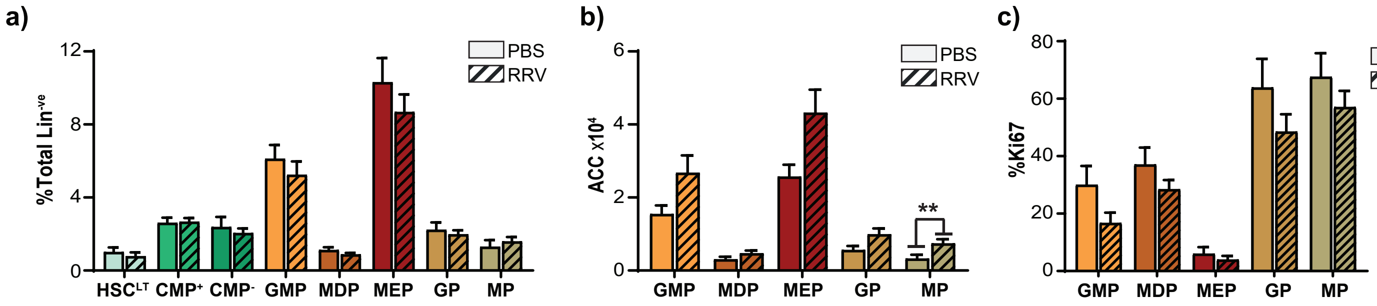


**Supplementary Figure 1. Perinatal liver inflammation leads to significantly increased number of myeloid progenitors with no changes in the percentage and proliferative capacity of downstream progenitor populations in PLI-recovered adult mice.** a) Bar graphs showing no change in the fraction of HSC$^{LT}$, CMP$^{+}$, CMP$^{-}$, GMP, MEP, GP and MP in the Lin$^{-ve}$ compartment in the bone marrow of PLI-recovered animals (n=22) c.t. PBS-injected controls (n=11). b) Bar graph showing quantification of ACC in GMP, MEP, GP, and MP with significant increase in MP in PLI-recovered animals (n=22) (p-value = 0.009+/- SEM) c.t. PBS-injected controls (n=11). c) Bar graph showing no change in Ki67% expression in GMP, MEP, GP and MP populations in PLI-recovered animals (n=22) c.t. PBS-injected controls (n=11). Error bars represent mean +/- SEM.

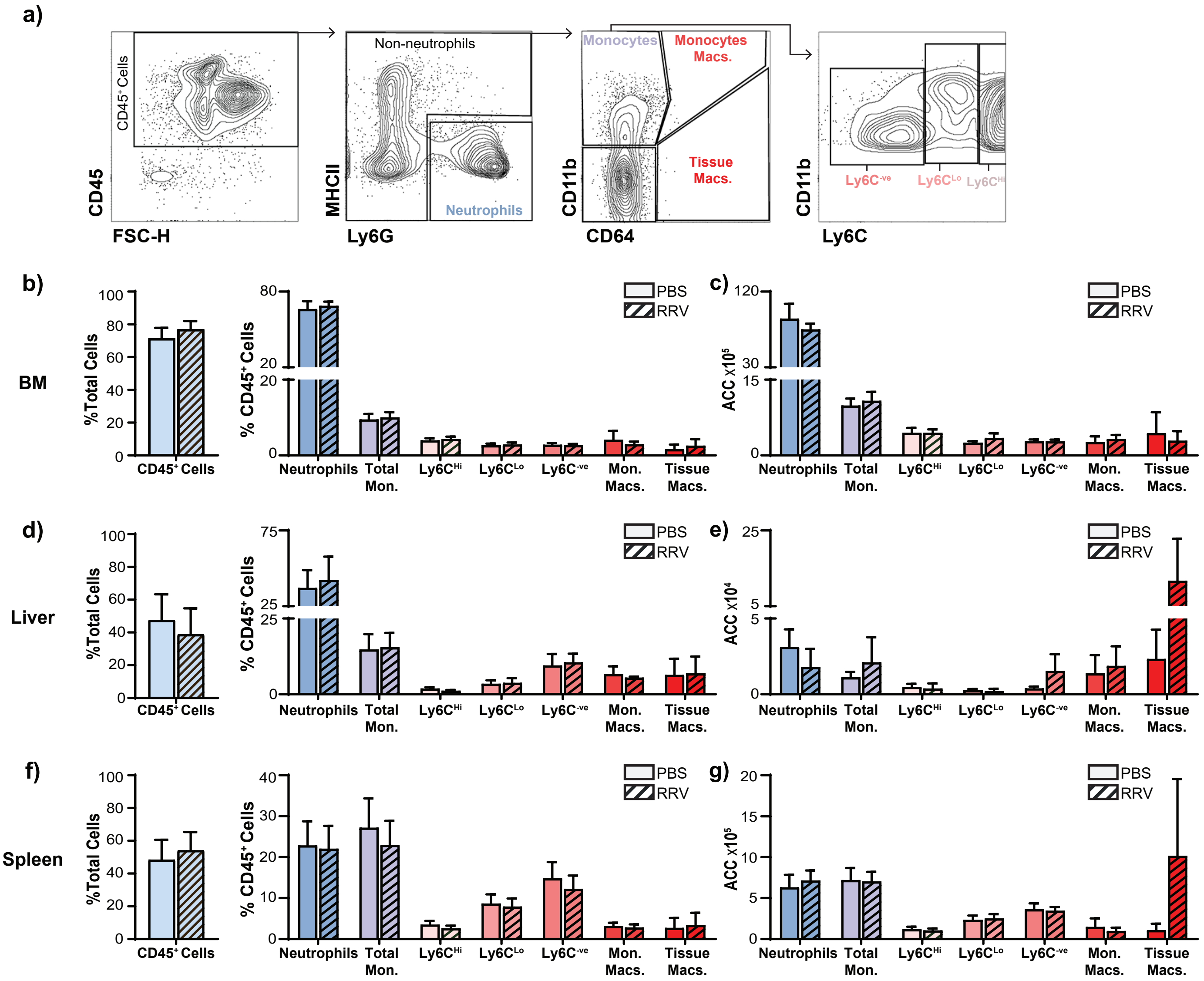


**Supplementary Figure 2. Perinatal liver inflammation does not lead to any quantitative changes in the mature myeloid compartment of PLI-recovered adult mice.** a) Plots demonstrating flow-cytometric gating strategy of mature myeloid populations in adult mice bone marrow. Bar graphs (left-right) showing quantification of percentage CD45$^+$ cells, percentage and ACC of mature myeloid populations in bone marrow (b and c), liver (d and e), and spleen (f and g) of PLI-recovered animals (BM: n=13, Liver: n=5, Spleen: n=14) c.t. PBS-injected controls (BM: n=12, Liver: n=6, Spleen: n=12). Error bars represent mean +/- SEM.
